# The role of vocalizations in agonistic interactions during competition for roosts in a solitary bat

**DOI:** 10.1101/2024.05.15.594314

**Authors:** Cristian Castillo-Salazar, Michael G. Schöner, Caroline R. Schöner, Gloriana Chaverri

## Abstract

Inter- and intraspecific competition for resources is common among individuals which share ecological niches. To avoid physical confrontations, individuals can use various types of signals to demonstrate their dominance, including vocalizations. *Kerivoula hardwickii* is a solitary bat species that lives in highly ephemeral plant structures, which are therefore a limited resource. So far, it is unknown if individuals of *K. hardwickii* use vocalizations during competitive encounters for roosts, and if the intention of these vocalizations can be deduced by potential rivals. We hypothesized that the calls emitted during roost competition contain information that influences the ability of an individual to defend its roost. We conducted roost competition experiments in a flight cage, where there was an individual roost owner and an intruder who would attempt to evict the owner from the roost. All the vocalizations emitted during these encounters were recorded and analyzed to determine which acoustic parameters, if any, had an influence on the successful defense of the roost. We found that the calls emitted by males can influence their ability to defend the roost, and that entropy is the parameter that most strongly explains a successful defense. High entropy suggests that encounters between individuals of *K. hardwickii* escalate to high levels of aggressiveness and explain whether calls influence an individual’s capacity to defend a roost. We suggest that bat vocalizations contain important information about individual characteristics, which in turn help bats make decisions during resource competition.

## Introduction

Competition for limited resources in the animal kingdom can escalate to physical contests that can lead to injury, increased energy expenditure, or even death (Briffa M & Elwood, 2004). Given the latter, most animals should prefer to resolve conflicts through signals instead of escalating to physical contests (Briffa, 2015). These signals may often demonstrate an animal’s fighting ability and allow it to gain or maintain its access to resources without battles (Bradbury & Vehrencamp, 2011). Some signals employed during contests can be color displays (Takeuchi, 2017), gestures (Call & Tomasello, 2020), facial expressions (Camerlink et al., 2018), body postures (Issa & Edwards, 2006), or movements (Reddon & Balshine, 2019). There are also chemical signals such as pheromones (Blaul & Ruther, 2012), tactile signals such as small physical contacts, and acoustic signals (Reichert & Gerhardt, 2013) that may be used during confrontations.

Acoustic signals are widely used for various activities related to competition or defense of resources (Reichert & Quinn, 2017). It has been documented that some types of calls may contain critical information about an individual’s competitive abilities, such as body size, aggressiveness, and even dominance rank in social species (Bradbury & Vehrencamp, 2011). These signals may also provide clues about a competitor’s defense potential or fighting motivation (Vannoni & McElligott, 2008). All this information provided by vocal signals may allow competitors to decide which individuals to engage in a fight (Ratcliffe et al., 2007). For example, male green frogs *Rana clamitans* (Anura, Ranidae) obtain information about an intruder’s body size from the frequency of its calls, so they alter their behavior according to the perceived size of intruders based on differences in the frequency spectra of acoustic signals (Bee et al., 2000). In free-ranging male baboons (*Papio cynocephalus ursinus*), the acoustic parameters of the calls they emit during interactions contain information about the male’s competitive ability. Dominant males emit higher fundamental frequencies and longer calls, while males that decrease in dominance emit shorter calls (Fischer et al., 2004). This use of acoustic signals to obtain information about individuals during competition has also been reported in other groups of mammals such as hyenas (Mathevon et al. 2010), deer (Reby & McComb 2003; Vannoni & McElligott, 2008), and rock hyraxes (Koren & Geffen, 2009). However, relatively few studies have been carried out on bats on this subject.

Bats are an ideal model for the study of acoustic communication in the context of social interactions given that sounds represent a vital signaling mode in this large mammalian taxon (Gillam & Fenton, 2016). Social calls in bats, for example, are known to contain information that is used in a wide variety of social contexts (Fernandez & Knörnschild, 2017; Jahelková et al., 2008). In bats that form groups, acoustic signals may facilitate the formation and maintenance of social groups (Chaverri & Gillam, 2016) and defense of communal roosts (Chaves-Ramírez et al., in press), while in solitary bats it has been documented that these calls are used to defend space and resources (Sun et al., 2021). Despite the latter, there are only a handful of studies that have addressed the role of social calls in solitary species and the contexts in which they are used, and particularly in the context of roost defense.

The woolly bat, *Kerivoula hardwickii* is a bat of the family Vespertilionidae found in Southeast Asia (Francis 2019). *K. hardwickii* is one of the few species of bats that are considered solitary because individuals do not join conspecifics while roosting. Individuals of this species have been found roosting in the pitchers of carnivorous plants of the genus *Nepenthes*, as well as developing tubular leaves of the families Zingiberaceae, Musaceae and Araceae (Grafe et al., 2011; McArthur 2012). Most plant structures used by *K. hardwickii* have a limited roosting space and usually only remain suitable for short periods of time (Schöner et al. 2017), so these types of roosts are relatively scarce resources. When resources are scarce it is expected that individuals should compete for their access and defend them (Grover et al., 1997). However, it is not known what acoustic signals can be emitted when several individuals attempt to secure access to a common roost. Therefore, the main goal of this study is to describe the acoustic signals emitted by individuals of *K. hardwickii* when they compete for a roost and to evaluate if the acoustic parameters of the calls have an influence on the ability of individuals to defend a roost. We hypothesized that the calls emitted during roost competition contain information that may help an individual to defend its roost (Reichert & Gerhardt, 2013), and that certain acoustic parameters potentially predict if an individual successfully accesses or defends the roost (Sun et. al, 2021).

## Methods

### Study Site

The study was conducted in Gunung Mulu National Park, located in Sarawak, Malaysia, from December 2017 to March 2018. This park has an extension of 544 km^2^ with altitudes that range from 50 to 2,376 masl. Gunung Mulu was declared a World Heritage Site by UNESCO in 2000 and it’s characterized by having a wide variety of vegetation types and abundance of karst areas (Proctor et al., 1983). Research permits were granted by the Sarawak Forestry Department Kuching, Sarawak, Malaysia (research permit no. NPW.907.4.4(JLD.14)-189 and park permit no. WL91/2017).

### Bat Sampling

Individuals were searched and captured in their roosts, the developing furled leaves of the plant families Zingiberaceae, Araceae, and Musaceae. We captured bats and placed them inside a cloth holding bag. For all the bats captured, we sexed and classified the reproductive status of the adult individuals. We determined age as adult or juvenile based on the level of ossification of the union of the epiphysis in the phalanges. For individual recognition, all adult males and non-reproductive females were marked with passive integrated transponders (ISO 11784/11785; Peddy-Mark, UK).

### Experiment

For the field experiments, a 3×3 m flight cage equipped with two video cameras (Sony HDR-CX560VE) was used. For sound recording, an Avisoft UltraSoundGate116Hn (Avisoft Bioacoustics, Glienike/Nordbahn, Germany) was used with an ultrasonic condenser microphone (CM16/CMPA, Avisoft Bioacoustics). Pregnant and lactating females were excluded from the experiments. All individuals selected for the experiments were returned to their original capture site in less than 24 hours. Water and insects, as a source of food, were provided *ad libitum* to individuals during and after the experiments.

To record the calls emitted during the interactions in which bats competed for a single roost, we conducted experiments inside a 3×3 m flight cage equipped with two video cameras (Sony HDR-CX560VE). First, a bat was released inside the flight cage until it entered and settled within the roost, a furled leaf of *Alpinia* sp. (Zingiberaceaea). The bat inside the roost was called the “owner”. Later, a second bat was released and all interactions between the two individuals were recorded. The flying bat was called the “intruder”. A successful defense of the roost was defined as when the owner bat was not evicted from the roost by the intruder within our experimental period of 30 min. A successful eviction was defined as instances when the intruder managed to evict the owner completely from the roost and did not allow it to return within a period of 30 minutes. The cameras and the microphone were placed as close as possible to the leaf to record all the interactions and calls that occurred inside the roost between the bats. Experiments were conducted only among same-sex individuals to avoid interactions between females and males that could not be directly associated with competition for roosts.

### Acoustic analysis

Acoustic analyzes were performed with the Avisoft-SASLab Pro version 5.1 software (Avisoft Bioacoustics, Glienike/Nordbahn, Germany). We generated spectrograms with an FFT length of 512 (frequency resolution = 1465 Hz) and 93.75% overlap (temporal resolution = 0.0427 ms). For each of the recorded calls, the acoustic parameters were measured in the maximum amplitude of the element and the mean spectrum of the entire element. The acoustic parameters duration, number of elements, element rate, peak frequency, minimum frequency, maximum frequency, bandwidth, and entropy, were estimated. The number of calls were counted and classified into categories according to the position of their tonal component (figure 1). We defined 5 categories, or types of calls, according to the position of its tonal component, where T1 corresponds to a call without a tonal component, T2 is a call with the tonal component at the beginning, T3 has a tonal component in the middle of the call, T4 is a call with the tonal component at the end, and T5 is a call which only consists of the tonal component. To prevent tonal components from altering the measurements of the acoustic parameters, we manually excluded them from the measured calls and all calls were measured as T1 type (Figure 1).

**Figure 1.**
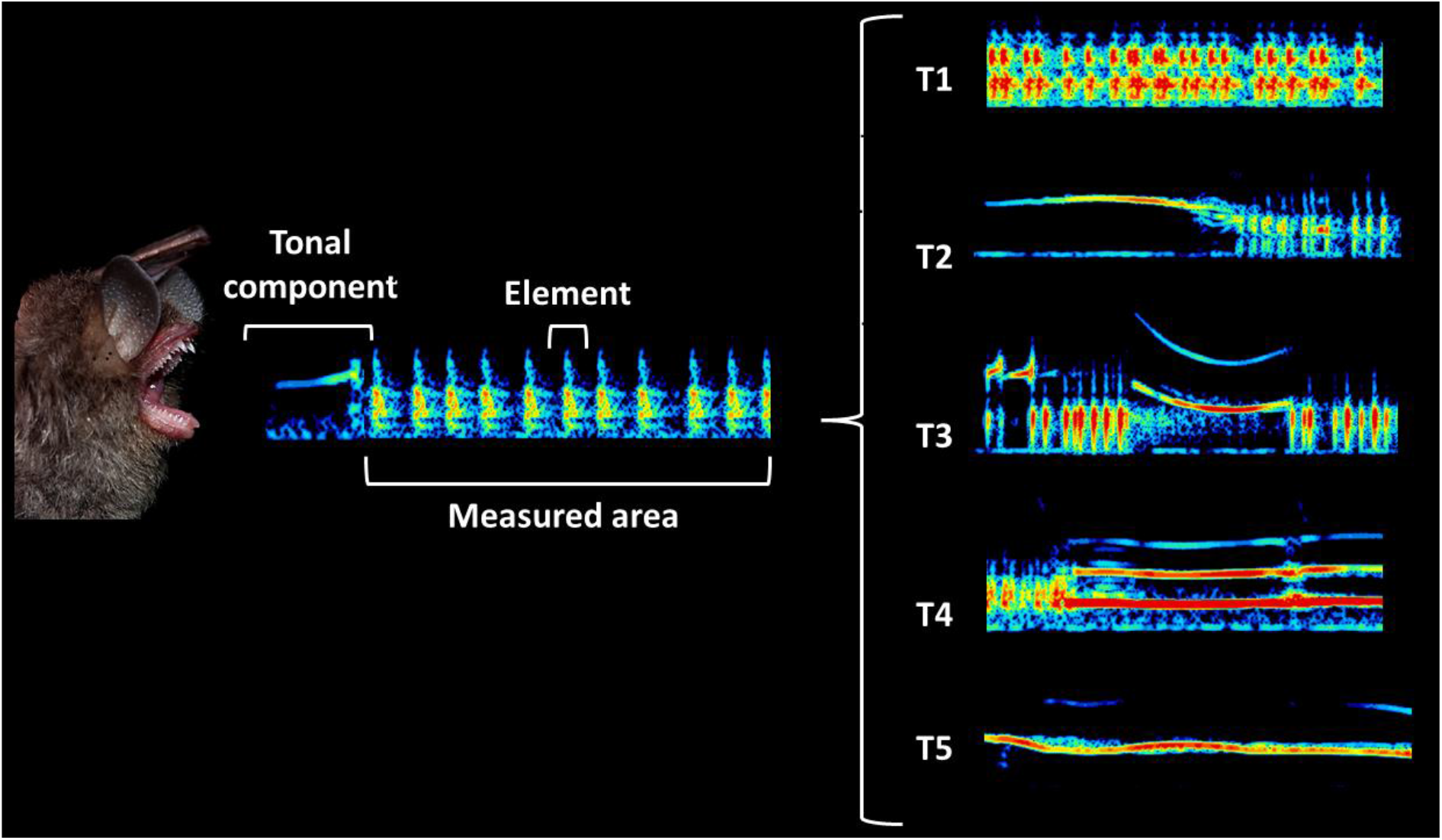
Classification of the calls according to the position of the tonal component emitted during the competition experiments for the roost in *K. hardwickii*.

### Statistical analysis

To determine which factors can predict roost defense (i.e., when the owner bat was not evicted from the roost by the intruder within our experimental period), we run generalized mixed models with a binomial distribution. First, to reduce the dimensionality between acoustic parameters and remove correlations between independent variables, we conducted a principal component analysis (PCA). Then, the components that explained the greatest variability among the acoustic parameters, which had an eigenvalue > 1, were extracted and used as independent variables in the models. The model ***Y(D) = m + S + P1 + P2 + P3 + P4 + I + O + error*** was used to determine which acoustic parameters of the calls could predict the successful defense of the refuge. Successful roost defense (D) was used as the dependent variable, sex (S) and extracted components (P1, P2, P3, P4) were used as the independent variables, while intruder ID (I) and owner ID (O) were considered random variables.

The model ***Y(D) = m + S+ NC + NT + I + O + error*** was used to determine if the number of calls and the number of types of calls could predict the successful defense of the roost. The successful defense of the refuge (D) was used as the dependent variable, sex (S), number of calls (NC) and number of types of calls (NT) as independent variables, while intruder ID (I) and owner ID (O) were used as the random variables. We performed the models for the complete data set first and then for the data separated by sex. The generalized mixed models were carried out with the lme4 package (Bates et al., 2015) and all statistical analysis were conducted in R v. 4.1.2 (R Core Development Team 2018).

## Results

To extract acoustic parameters, 259 calls corresponding to 34 pairs of eviction experiments were analyzed, 31 experimental pairs were female and 24 pairs were male; 67 calls were emitted by female pairs, and 194 calls were emitted by male pairs. From the PCA analysis, 4 components were extracted with eigenvalues greater than 1. These 4 components explained 75.4% of the variance. The first component explains 29.2% of the variance and is related to the maximum frequency and bandwidth parameters. The second component explains 21.5% of the variance and is related to entropy. The third component explains 15.9% of the variance and is related to the peak frequency, while the fourth component explains 8.6% of the variance and is related to the duration of the call (Table 1).

**Table 1.** Loadings for the four components extracted of the PCA from the roost eviction experiments, with the proportion of variation explained by each component.

| Parameter | Components |  |  |  |
| --- | --- | --- | --- | --- |
|  | PC1 | PC2 | PC3 | PC4 |
| Duration | 0.073327 | 0.050984 | 0.21927 | **0.730061** |
| Element rate | 0.062051 | 0.293912 | -0.2927 | 0.17166 |
| Pf<sub>max</sub> | 0.011846 | 0.289882 | **0.505527** | -0.25248 |
| Minf<sub>max</sub> | 0.181909 | 0.271237 | 0.396006 | 0.317312 |
| Maxf<sub>max</sub> | **-0.44993** | 0.219253 | 0.13715 | -0.04179 |
| Bandw<sub>max</sub> | **-0.48143** | 0.058127 | -0.0827 | -0.19511 |
| Entropy<sub>max</sub> | -0.01189 | **0.487469** | -0.34855 | -0.11885 |
| Pf<sub>mean</sub> | 0.087766 | 0.257223 | **0.457404** | -0.34044 |
| Minf<sub>mean</sub> | 0.203065 | 0.392958 | -0.0615 | -0.06676 |
| Maxf<sub>mean</sub> | **-0.48016** | 0.068954 | 0.095332 | 0.187984 |
| Bandw<sub>mean</sub> | **-0.49263** | -0.00798 | 0.098966 | 0.191129 |
| Entropy<sub>mean</sub> | -0.03174 | **0.488815** | -0.27359 | 0.164169 |
| Variation% | 29.287 | 21.573 | 15.914 | 8.638 |

Bold values indicate the highest loading associated with a particular component.

In the mixed model of both sexes (AIC = 47.589), which estimates the contribution of various acoustic parameters (based on the 4 principal components) to the defense of the roost, we found that sex, PC2 and PC3 are the variables that explain roost defense (Table 2). Therefore, the data were separated by sex to determine how acoustic parameters explained roost defense while removing the effect of sex. For females alone, we found that none of the acoustic parameters explained roost defense. For males, PC2 (i.e., entropy; Table 1) was significantly and positively associated with roost defense (Figure 2). The number and the types of calls did not seem to influence the defense of the roost (Table 2).

**Table 2.**
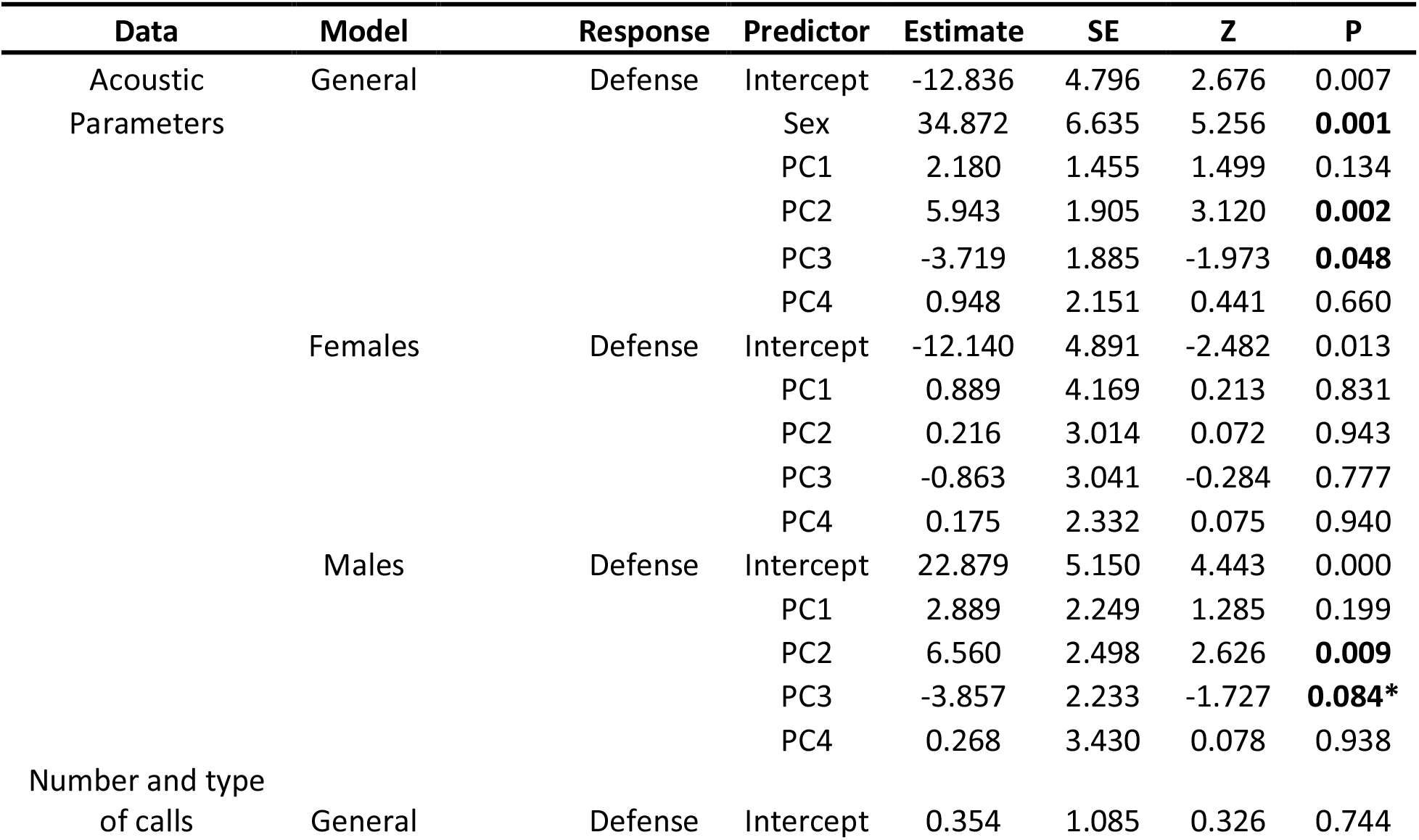
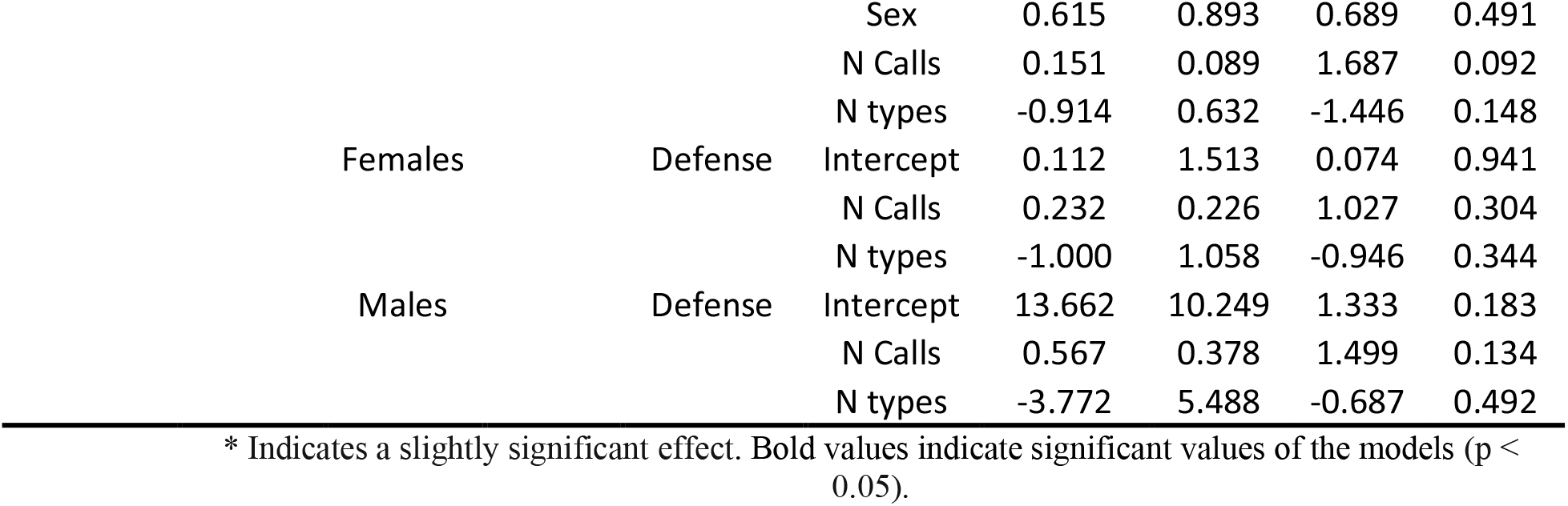
Mixed generalized linear models between acoustic parameters and complexity of calls on successful roost defense.

**Figure 2.**
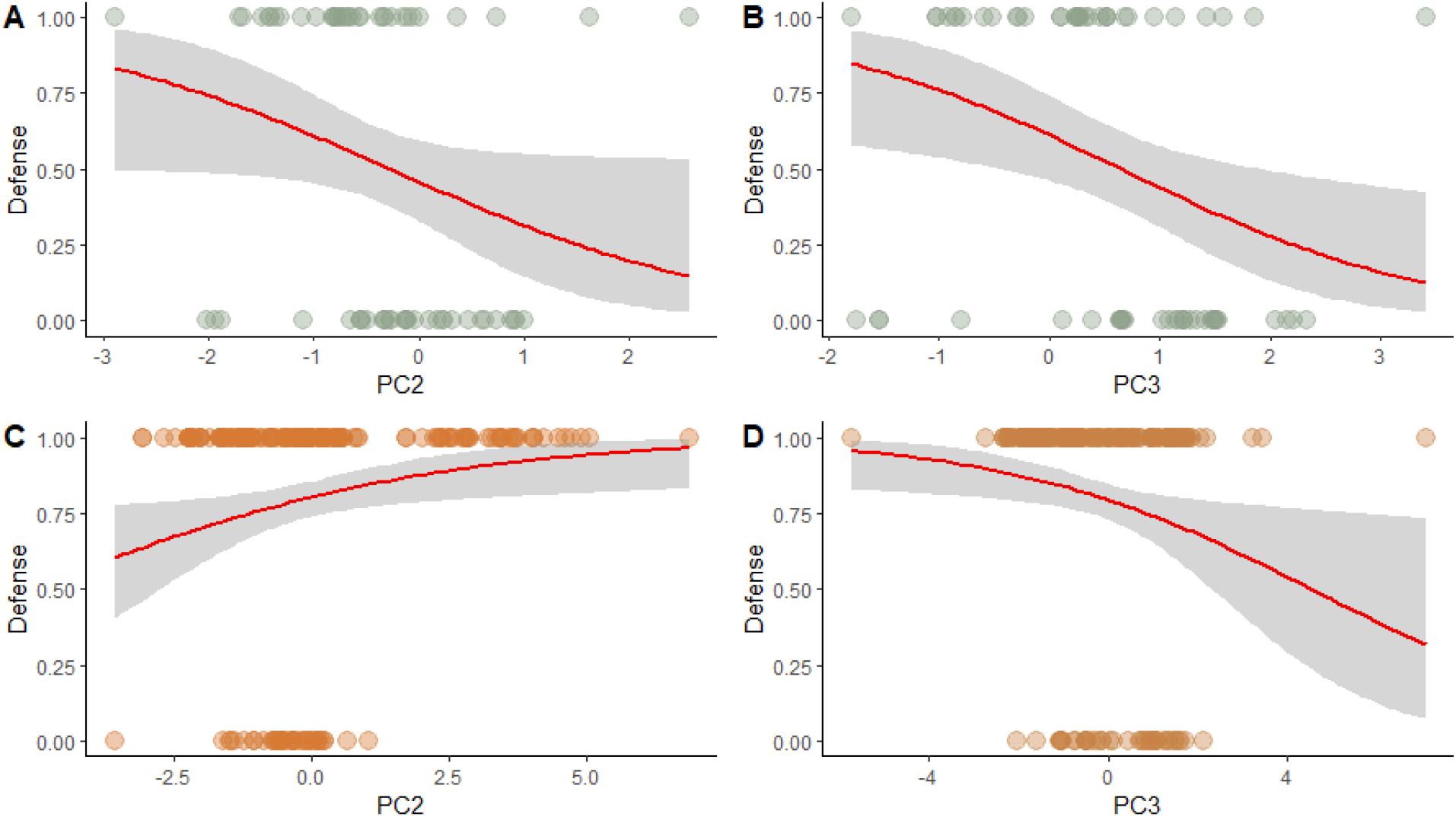
Mixed generalized linear models between acoustic parameters and successful roost defense (denoted by y-values = 1). Models A and B correspond to the effect of the variables PC2 and PC3 on the defense of the roost in experiments that only include females. Models C and D correspond to the effect of the variables PC2 and PC3 on the defense of roosts in male-only experiments.

## Discussion

We found that the calls emitted by male individuals of *K. hardwickii* during their interactions may predict their ability to defend their roost sites. Entropy was the acoustic parameter that most strongly explained successful roost defense. These results support our hypothesis that calls emitted during roost competition contain information that may influence the result of dyadic contests in *K. hardwickii*.

The level of aggressiveness in the calls of several species has often been related to their frequency and complexity. For example, in the gray tree frog *Hyla versicolor*, males that won aggressive interactions tended to have lower-frequency aggressive calls than losers (Reichert & Gerhardt, 2013). In bats, it has been found that in the Great Himalayan leaf-nosed bat (*Hipposideros armiger*) the peak frequency of territorial calls indicates the dominance of the individual in the colony (Sun et. al, 2021). However, in *K. hardwickii* the parameters related to a call’s frequency did not predict roost defense; rather, it was the call’s entropy that seemed to favor roost defense. Basically, there was a greater probability of roost defense when the bat within the roost emitted calls with higher entropy.

Noisy, or high entropy, vocalizations have been documented to be related to the degree of arousal and alertness (Fitch et al. 2002). The latter states cause changes in physiology, such as increased respiration and heart rate, changes in the pressure of the glottis, and changes in the phonation of the call (Berry et al., 1996). The increase in pressure flow in the glottis generates calls with higher entropy and produce changes in the vocal structure of the individual (Goudbeek & Scherer, 2010; Lemasson et al., 2012). For example, it has been documented that in some primate species, such as Geoffroy’s Spider Monkeys (*Ateles geoffroyi*) and rhesus macaques (*Macaca mulatta*), there is a transition from a low-entropy vocalization to a high-entropy vocalization when the aggression changed from low severity to high severity (Fitch et al. 2002; Ordóñez-Gómez et al., 2015). It has also been documented that the Java sparrow (*Lonchura oryzivora*) produces similar calls in different situations of aggressiveness or affinity, but entropy and sound pressure levels were higher in aggressive calls than in affiliative calls (Furutani et al., 2018). This indicates that entropy increases considerably when the severity of the encounter increases, and some research suggests that high-entropy vocalizations can produce negative responses in the mammalian auditory system and aim to discourage an opponent’s aggressiveness (Gouzoules & Gouzoules, 2000). Documented calls with high entropy during encounters suggest that male *K. hardwickii* reach levels of high severity and aggressiveness during competitive interactions. Basically, our results suggest that as the entropy of the call emitted by the defending individual inside the roost increases, it is more likely that this individual will repel an intruder.

Our study is, to the best of our knowledge, the first to describe aggressive competitive interactions in males of a strictly solitary bat. The few published studies on direct competition and aggressive interactions between bats have been carried out with species that live in colonies (Chaves-Ramírez et al., in press; Fernandez et al., 2014). In these groups, the bats have constant interactions with individuals of the colony, which generates dominance hierarchies in the group (Zhao et al., 2018). Calls emitted during aggressive interactions in these colonies have been shown to contain information about body size, dominance rank, and individual identity (Sun et al., 2021). In our case, *K. hardwickii* is a bat that roosts alone and does not form colonies. In addition, a previous study (C. Castillo-Salazar, unpublished data) showed that body mass did not influence aggressiveness or the eviction of individuals from roosts. Therefore, we suggest that the calls can have information about individual identity and levels of aggressiveness that help individuals to gauge each other’s abilities during competition for roosting resources.

## Acknowledgement

We want to give a special thanks to Nikolaj Meyer, Johanna Lauffer, Hanna Halblang, Lioba Uffer, Sofie Gaw, Jon A. Romero, Julien Bota, Chai Shong Kian and Aubrey Siebels for all the help in the field and the great company during the months of the study. Thanks to Ellen McArthur and Faisal Ali for helping us with all the paperwork and letters so that the investigation could be carried out. Thanks so much to all the family of Mulu National Park who welcomed us warmly and made our stay very special.

